# Investigating Combined Hypoxia and Stemness Indices for Prognostic Transcripts in Gastric Cancer: Machine Learning and Network Analysis Approaches

**DOI:** 10.1101/2024.06.26.600775

**Authors:** Sharareh Mahmoudian-Hamedani, Maryam Lotfi-Shahreza, Parvaneh Nikpour

**Author notes:** Correspondence: Parvaneh Nikpour (PhD), Department of Genetics and Molecular Biology, Faculty of Medicine, Isfahan University of Medical Sciences, Isfahan, Iran. and.

## Abstract

**Introduction:** Gastric cancer (GC) is among the deadliest malignancies globally, characterized by hypoxia-driven pathways that promote cancer progression, including mechanisms associated with stemness facilitating invasion and metastasis. This study aimed to develop a prognostic decision tree using genes implicated in hypoxia and stemness pathways to predict outcomes in GC patients.

**Material and Methods:** GC RNA-seq data from The Cancer Genome Atlas (TCGA) were utilized to compute hypoxia and stemness scores via Gene Set Variation Analysis (GSVA) and mRNA expression-based stemness index (mRNAsi). Hierarchical clustering based on these scores identified clusters with distinct survival outcomes, and differentially expressed genes (DEGs) between these clusters were identified. Weighted Gene Co-expression Network Analysis (WGCNA) was employed to identify modules and hub genes associated with clinical traits. Hub genes overlapping with DEGs were extracted, followed by functional enrichment, protein-protein interaction (PPI) network analysis, and survival analysis of shared genes. A prognostic decision tree was constructed using survival-associated genes.

**Results:** Hierarchical clustering identified six clusters among 375 TCGA GC patients, showing significant differences in survival outcomes between cluster 1 (with low hypoxia and high stemness) and cluster 4 (high hypoxia and stemness). Validation in the GSE62254 dataset corroborated these findings. WGCNA revealed modules correlating with clinical traits and survival. Functional enrichment highlighted pathways such as cell adhesion and calcium signaling. The decision tree based on survival-related genes including *AKAP6*, *GLRB*, *LINC00578*, *LINC00968*, *MIR145*, *NBEA*, *NEGR1* and *RUNX1T1* and achieved an area under the curve (AUC) of 0.81 (training) and 0.67 (test), demonstrating the utility of combined scores in patient stratification.

**Conclusion:** This study introduces a novel hypoxia-stemness-based prognostic decision tree for GC. The identified genes show promise as prognostic biomarkers for GC, warranting further validation in clinical settings.

## Introduction

Gastric cancer (GC) is the third cause of global cancer-related fatalities and is responsible for approximately one out of every twelve oncological deaths. It is also the fifth most frequently occurring form of cancer, accounting for 5.7% of all new cases (1). Despite a decrease in incidence rates, which is a result of better food preservation practices and improved living conditions associated with economic development, increasing the aging population will lead to a rise in GC cases in the future (2,3).

Intratumor heterogeneity is a major attribute of GC which contributes to negative outcomes such as tumor relapse and chemo resistance (4). One of the factors associated with heterogeneity is hypoxia; higher hypoxia is linked with augmented mutational load across various cancer types (5). Oxygen serves as an electron acceptor in numerous organic and inorganic reactions and thus, is considered a critical element of aerobic metabolism. Hypoxia, a condition in which the amount of Oxygen in the tissues is less than 2% (6), promotes aggressive and metastatic cancer by enhancing tumor cell proliferation, survival, immune evasion, inflammation, angiogenesis, and invasion (7). Hypoxia can lead to the activation of hypoxia-inducible factors (HIFs), a group of transcription factors that are engaged in the regulation of various genes involved in cancer biology, particularly those required in stem cell specification, maintenance, and survival (8). Cancer stem cells account for a small number of cancer cells with stem cell attributes, namely self-renewal and differentiation into other cancer cells (9). Stemness is characteristic of the most lethal cancers, like lung cancer, breast cancer, colorectal cancer and leukemia (10).

Numerous investigations have been conducted regarding the hand-in-hand effects of hypoxia and stemness in cancer progression. For example, Zhao et al showed that chemical induction of hypoxia results in an increase in lung cancer cell stemness and drug resistance in CD166-positive stem cells. Of note, patients with higher expression of CD166 showed poorer prognoses as well (11).

It is also demonstrated that post-hypoxic breast cancer cells with the ability to metastasize, exhibit an enrichment in pathways that are associated with the expression of cancer stem cell-related genes. These cells are less sensitive to chemotherapeutics and lead to recurrence after treatment (12).

In the past decade, The Cancer Genome Atlas (TCGA) has provided a detailed understanding of primary tumor landscapes through the generation of comprehensive multi-omics characteristics and annotations of pathophysiological features and clinical information (13). Recently, many studies have employed TCGA data along with bioinformatics tools and machine learning algorithms to provide insight into underlying cellular mechanisms of GC and suggest tools to predict the patients’ prognoses, but little investigation has been carried out on the possibility of suggesting a machine learning model based on hypoxia and stemness indices to predict the prognosis of GC patients.

In the present study, RNA-seq data of stomach adenocarcinoma (STAD) patients were downloaded from TCGA and normalized using the edgeR package. Hypoxia and stemness scores of the samples were calculated using GSVA (gene set variation analysis) package and mRNAsi (mRNA stemness index) methods in R, respectively. Next, based on these scores, two-dimensional hierarchical clustering and survival analysis were performed on TCGA-STAD samples and differentially expressed genes (DEGs) were then identified between the two clusters exhibiting the most significant variations in survival rates. Meanwhile, WGCNA (weighted gene co-expression network analysis) was carried out and hub genes of the survival-related clusters were selected. The shared genes between the WGCNA hub genes and the DEGs were then identified. Survival analysis was performed on these shared genes, and using the genes that were significantly associated with survival, a prognostic decision tree model was constructed and the related AUCs were calculated. The flowchart of the current study is presented in Figure 1.

**Figure 1.**
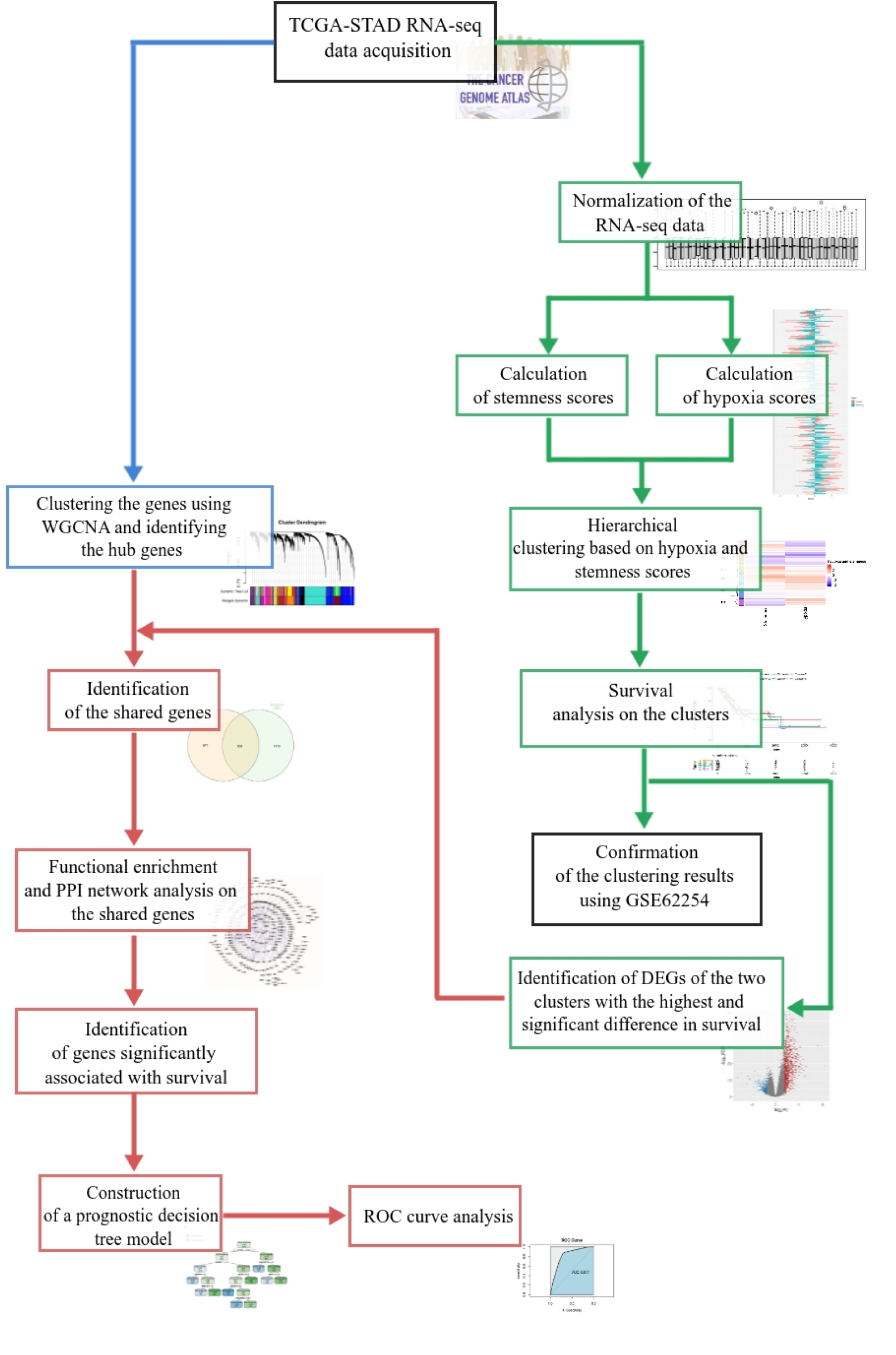
The workflow of the present study. In the workflow above, the study is divided into three parts that are identified with different colors, the first part of the study includes clustering the TCGA-STAD RNA-seq data based on hypoxia and stemness scores (green boxes and arrows), the second part includes WGCNA and identifying the hub genes (the blue box and arrow) and the third part includes identification of the shared genes between the first and the second parts and constructing a decision tree model using the survival-related shared genes (pink boxes and arrows). DEGs: Differentially-expressed genes, ROC: receiver operating characteristic, STAD: Stomach adenocarcinoma, TCGA: The Cancer Genome Atlas, WGCNA: Weighted gene co-expression network analysis,

## Materials and methods

### Data acquisition and processing

RNA-seq data of 375 STAD samples (tumoral) and related clinical data were retrieved from TCGA using the TCGAbiolinks R package (14). The RNA-seq data in the form of STAR (Spliced Transcripts Alignment to a Reference) counts were normalized using the trimmed mean of M values (TMM) method and log_2_-transformed using the edgeR package (15). As a validation set, the microarray gene expression data of 300 GC patients (GSE62254 (16)) were downloaded from Gene Expression Omnibus (GEO) database via the GEOquery package (17) and normalized using the limma (18) package. The related clinical data of GSE62254 were retrieved as well (16).

### Calculation of hypoxia score using GSVA

HALLMARK_HYPOXIA.v7.5.1, a gene set containing 200 hypoxia-related genes was retrieved from the Molecular Signatures Database (MSigDB, https://www.gsea-msigdb.org/gsea/msigdb), which provides a vast collection of gene sets with annotations that can be utilized to detect patterns of different pathways using samples’ gene expression data. Hypoxia scores of TCGA-STAD and GSE62254 samples were subsequently estimated employing the GSVA R package by using the HALLMARK_HYPOXIA.v7.5.1 as the gene signature and ssGSEA as the method.

### Calculation of stemness scores using mRNAsi method

In 2018, Malta et al proposed a method to calculate the mRNA expression-based stemness index (mRNAsi) of cancer cells, using the one-class logistic regression (OCLR) machine-learning algorithm (19,20). In the present study, mRNAsi was calculated by correlation analysis between stemness signatures’ weight vector and mRNA expression in GC samples obtained from TCGA-STAD and GEO database (GSE62254) was calculated using Spearman’s method.

### Unsupervised two-dimensional hierarchical clustering based on hypoxia scores and mRNAsi of GC samples

To identify and confirm the association between hypoxia and stemness scores and prognosis of GC patients, first an unsupervised two-dimensional hierarchical clustering based on these two scores was carried out on the TCGA-STAD and GES62254 samples. The results of the hypoxia score and mRNAsi of GC samples were first standardized using the “scale” function in the R. To perform hierarchical clustering, the Manhattan method for distance measurement and Ward’s minimum-variance method for linkage analysis were employed and the number of clusters was set to six (k=6). Distance measurement and linkage methods are used to calculate the closeness of the samples and the distance of the clusters, respectively (21). The same methods were applied to the results of hypoxia scores and mRNAsi of GC samples of the GSE62254 dataset (As the validation set). The clustering was performed and visualized using the hclust and ComplexHeatmap (22) packages, respectively.

### Survival analysis and identification of DEGs of GC samples of TCGA-STAD

Survival analysis was performed on the clustering results of TCGA-STAD and GSE62254 validation set to unravel the association of hypoxia and stemness-based hierarchical clustering with prognosis and possible pattern of hypoxia and stemness in groups with high and low survival rates. To this aim, the survival package in R was employed. The overall survival (OS) of distinct clusters was evaluated using the Kaplan-Meier curve and log-rank test. Next, DEGs of the two TCGA-STAD derived clusters with the highest difference in survival rate were identified using false discovery rate (FDR)<0.05 and |log_2_ fold change (log_2_ FC) |>1.5 by employing the edgeR package in R.

### Weighted gene co-expression network analysis

WGCNA is a systems biology analysis method for categorizing the genes in the microarray/RNA-seq samples into distinct modules based on their correlation. It can also be utilized for associating modules with samples traits in order to find the most pertinent modules for targeted therapy or introducing potential biomarkers (23). To construct a weighted gene co-expression network, first TCGA-STAD samples were clustered using the average method. Next, an adjacency matrix was formed as a result of converting the similarity matrix using the β value as the soft threshold. To create a scale-free network, a topological overlap matrix (TOM) was then created and by applying DynamicTreeCut method, TOM was converted into a weighted gene co-expression network and each module was identified by a specific color. To correlate each module with different clinical traits, module eigengenes were calculated. Module eigengenes are described as the first principal components that summarize the overall gene expression level in individual modules. After selecting those modules that were correlated with clinical traits related to GC patients’ prognosis, the module membership (MM) ≥ 0.7 was used to identify hub genes which were associated with prognosis. The higher value of GS represents the more prognostic value it holds for the patient.

### Functional enrichment and protein-protein interaction (PPI) network analyses

To identify the shared genes between the previously mentioned DEGs and hub genes attained from WGCNA, InteractiVenn (24), an online tool for creating Venn diagrams was employed. The Kyoto Encyclopedia of Genes and Genomes (KEGG) and Gene Ontology (GO) terms associated with the shared genes were then obtained from the Database for Annotation, Visualization and Integrated Discovery (DAVID) (25) and visualized using the ggplot2 package in R. To construct a PPI network, list of shared genes was applied to STRING database and the network was constructed with minimum interaction score of 0.7 (high confidence) (26), and the resulted network was exported to Cytoscape software (27).

### Survival analysis for shared hub genes and DEGs

To explore the association of any of the shared genes with overall survival, the survival package was employed in R. Statistical significance of the association was calculated using the log-rank test and *p*. value < 0.05 was considered as the threshold.

### Creating a machine learning-based prognostic decision tree

A prognostic decision tree model was constructed using the survival-related shared genes as inputs and the Recursive Partitioning and Regression Trees (RPART) algorithm. The RPART algorithm which is implemented in rpart R package (28), is a method for building classification and regression trees that can handle both categorical and continuous variables, missing values, and complex interactions. The data was split into a training (75%) and a testing (25%) set and the model was fitted on the training set. The rpart.plot package in R (29) was used to visualize the decision tree. The performance of the model was then evaluated on the testing set by plotting the receiver operating characteristic (ROC) curve and calculating the area under the curve (AUC) using the pROC package in R.

## Results

### Association between the combined hypoxia and stemness with GC patients’ survival

To discover the correlation between combined hypoxia and stemness and the survival of GC patients, an unsupervised two-dimensional hierarchical clustering was performed based on the combined hypoxia and stemness scores of 375 GC patients whose tumoral samples’ transcriptomic data was retrieved from TCGA. The result of clustering is represented in figure 2a which included: cluster 1 (66 samples, low hypoxia and high stemness), cluster 2 (45 samples, low hypoxia and stemness), cluster 3 (84 samples, rather high hypoxia and high stemness), cluster 4 (53 samples, high hypoxia and stemness), cluster 5 (80 samples, rather high hypoxia and rather low stemness) and cluster 6 (44 samples, low hypoxia and stemness). According to the survival analysis performed on these clusters, the highest survival difference was attributed to clusters 1 (higher survival rates) and 4 (lower survival rates), and log-rank test result trended towards significance (*P .*value =0.06, Figure 2b). Then, we determined whether we could identify the previously described hypoxia-stemness clusters in an external validation dataset, GSE62254. We applied the same hierarchical-clustering algorithm to subgroup the GC patients according to the combined hypoxia and stemness scores (Figure 2c) and survival rates were then calculated for each cluster. Our results showed that the difference in survival rate between the two clusters with low hypoxia and high stemness (cluster 4 with higher survival rates) and high hypoxia and stemness (cluster 6 with lower survival rates) was statistically significant (*P .*value =0.01, Figure 2d). These results confirmed the accuracy of the combined hypoxia and stemness scores in determining patient risk stratification and prognosis. After confirming the prognostic efficacy of the combined scores, we extracted 1446 differentially-expressed genes between clusters 1 (higher survival rates) and 4 (lower survival rates) of the TCGA-STAD samples.

**Figure 2.**
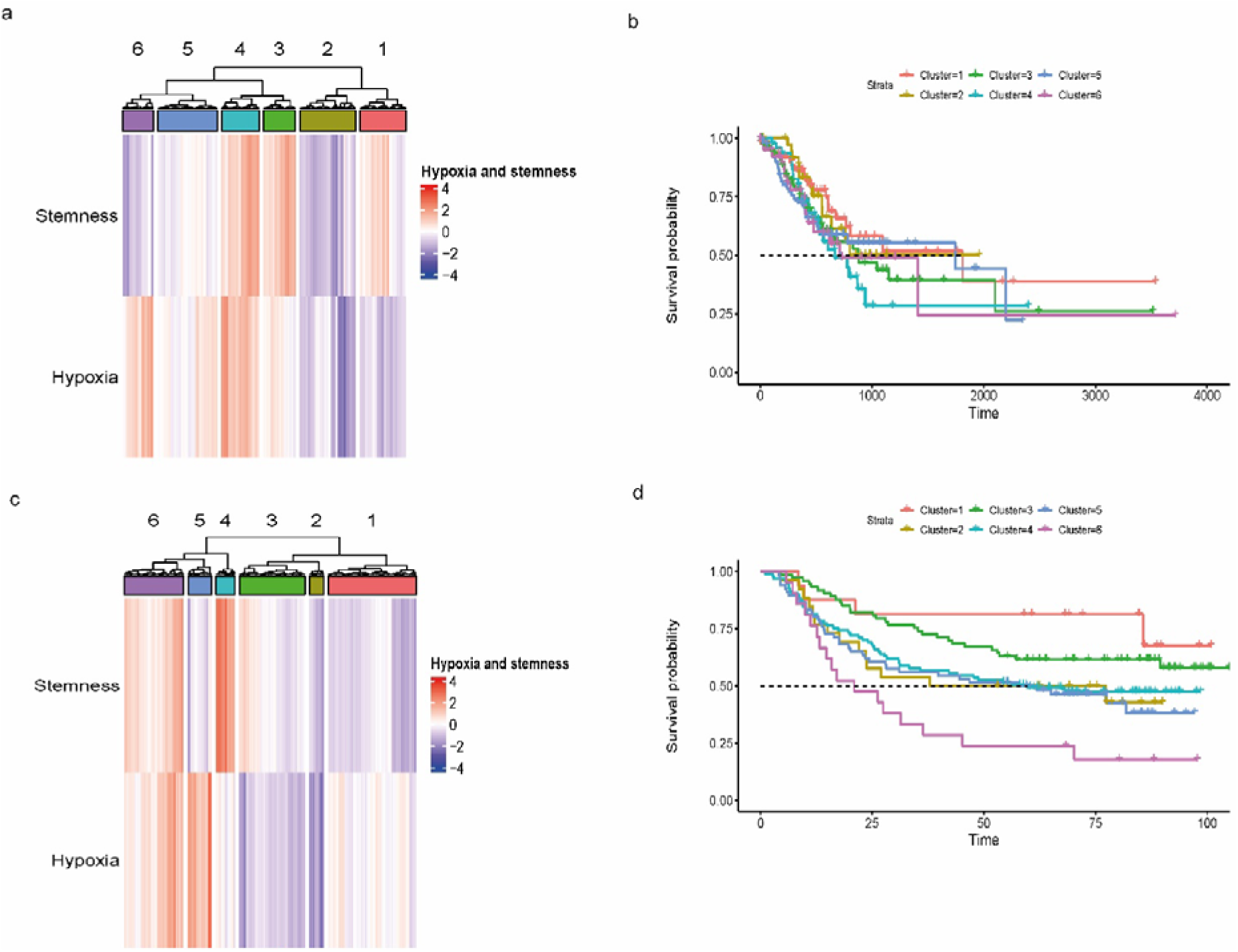
Clustering and survival analysis of TCGA-STAD and GSE62254 datasets. TCGA-STAD and GSE62254 dataset samples were clustered based on hypoxia and stemness scores (a and c), then survival analysis were performed for the clusters and Kaplan-Meier plots were extracted (b and d). STAD: Stomach adenocarcinoma, TCGA: The Cancer Genome Atlas

### Identification of prognosis-related modules using WGCNA

To define the prognosis-related key gene clusters and hub genes of GC, we performed WGCNA analysis using the normalized transcriptomic data of 375 GC samples retrieved from TCGA and the matched clinical data including pathologic stage, pathologic_T, pathologic_N, pathologic_M, days to death and overall survival. The soft threshold was set to 9, and the scale-free topology fitting index reached 0.89 (Figures 3a and 3b). Thirty modules were then detected using the WGCNA calculation which were merged into 15, each with a unique color, by a cut-off of 0.4, while the minimum number of genes in each cluster was set to 30 (Figure 3c). The genes contained in the dark gray module are genes that do not belong to any other modules. Pearson correlation analysis was employed to identify modules that were significantly linked to the prognosis-related clinical traits. There found to be 15 modules, of which dark turquoise (R=0.16, *P*. value =0.04), dark red (R=0.15, *P*. value =0.003), and dark orange (R=-0.15, *P*. value =0.005) were related to the pathologic stage. Blue (R=0.15, *P*. value =0.004), dark turquoise (R=0.19, *P*. value =2e-04), dark red (R=0.17, *P*. value =0.001), dark green (R=-0.14, *P*. value =0.006), dark orange (R=-0.2, *P*. value =1e-04) and royal blue (R=-0.16, *P*. value =0.001) were related to pathologic_T, magenta (R=0.12, *P*. value =0.02) and dark orange (R=-0.16, *P*. value =0.002) were related to pathologic_N and greenyellow (R=0.16, *P*. value =0.002) and purple (R=0.11, *P*. value =0.03) modules were related to overall survival (Figure 3d). With module membership (MM)>= 0.7 being set as a threshold to select hub genes, there were overall 1885 genes selected for further analysis. These genes were then aligned to the differentially-expressed genes between clusters 1 and 4 of the TCGA-STAD samples and 569 final shared genes, which henceforth will be addressed as the “**shared genes**” were extracted for further analysis (Figure 4).

**Figure 3.**
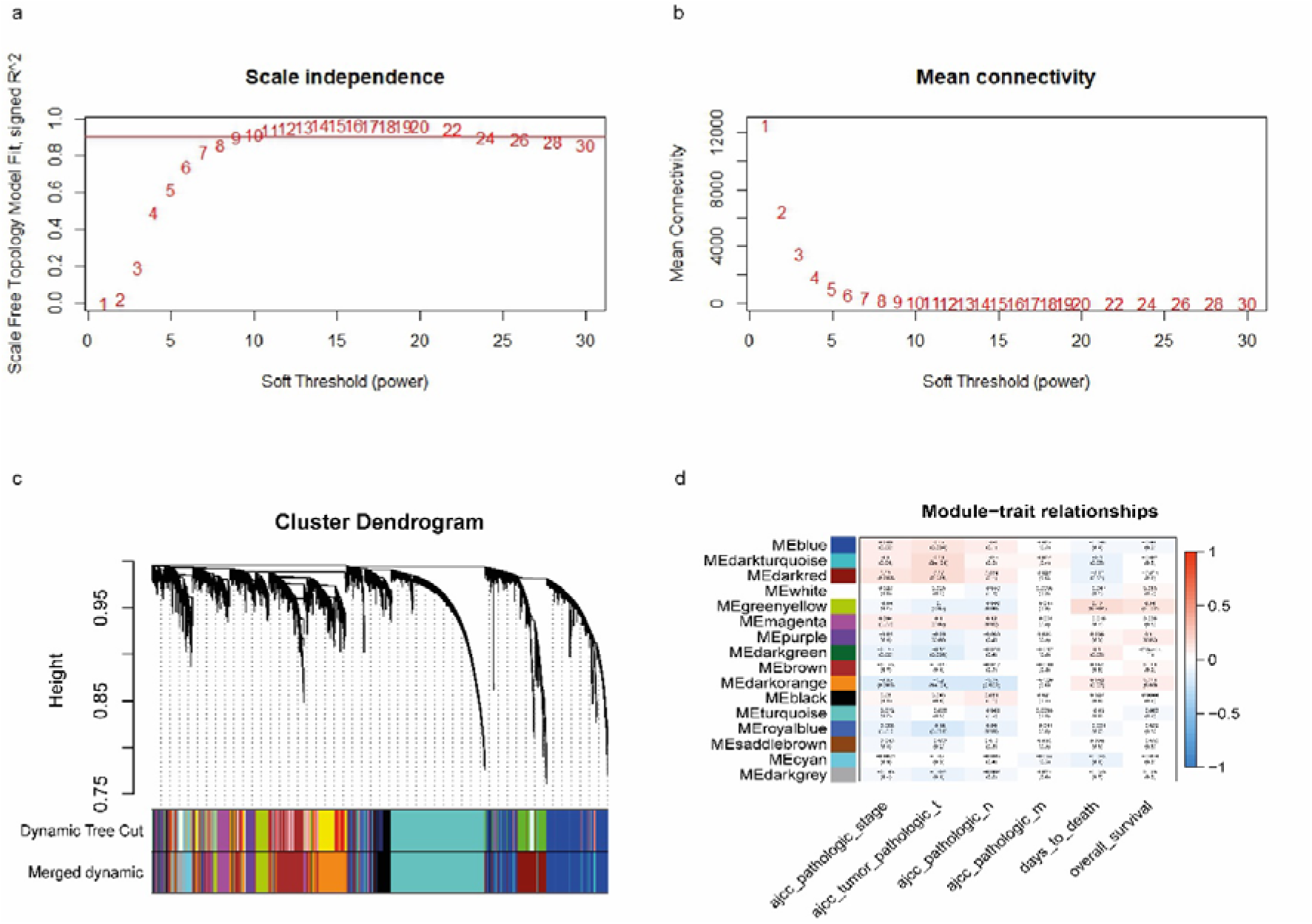
The WGCNA results. The plots represent scale-free fit index versus soft-thresholding power (a) and mean connectivity versus soft-thresholding power (b), respectively. The cluster dendrogram of gene expression data shows distinct modules with different colours representing each one (c). Module-trait relationships has been represented as a heatmap (d). ME: Module characteristic gene, WGCNA: Weighted gene co-expression network analysis

**Figure 4.**
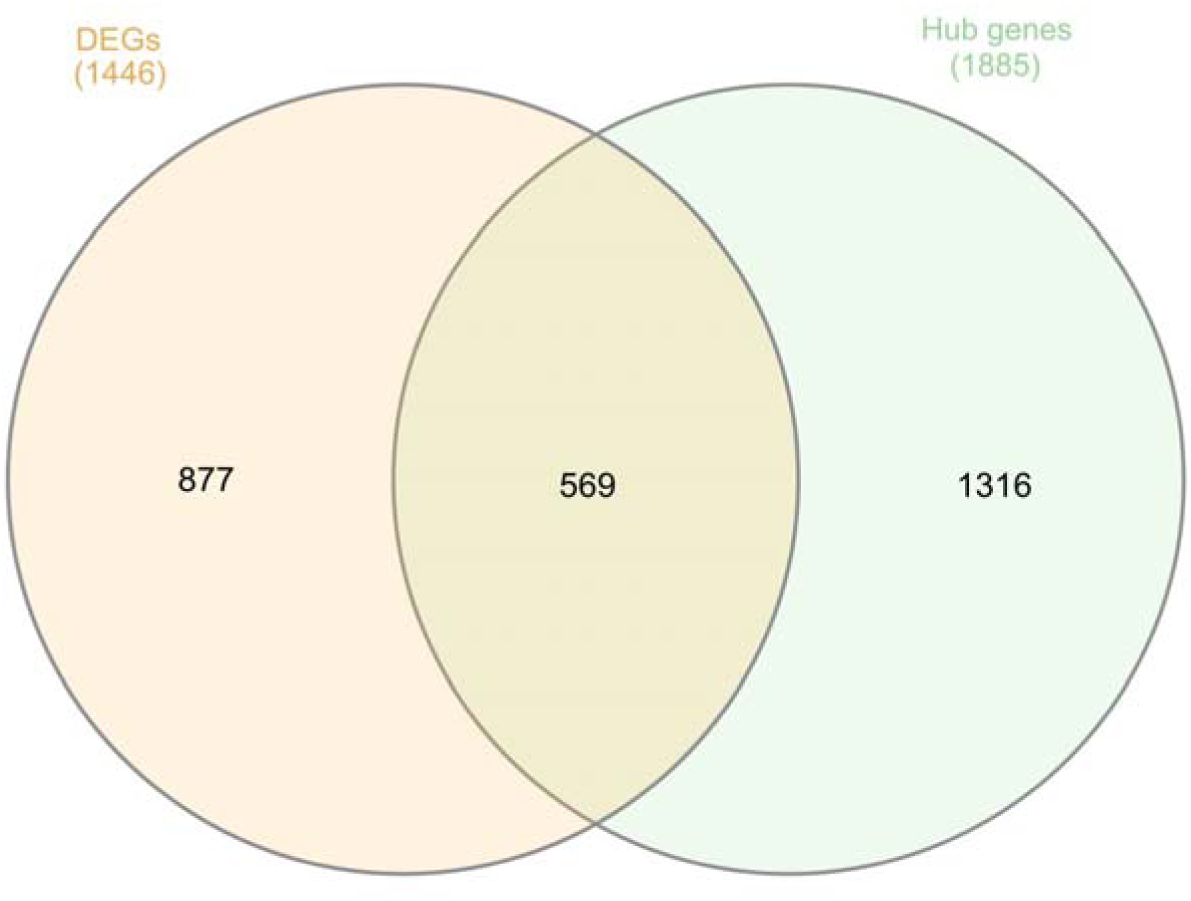
Venn diagram representing the shared genes between the hub genes and the DEGs. The Venn diagram shows the number of shared genes (#596) between the hub genes resulted from WGCNA (#1885) and differential expression analysis of cluster 1 *vs.* 4 of TCGA-STAD clustering results (#1446). DEGs: Differentially-expressed genes, STAD: Stomach adenocarcinoma, TCGA: The Cancer Genome Atlas, WGCNA: Weighted gene co-expression network analysis.

### Functional enrichment analysis and PPI network of the shared genes

In order to gain further insight into the functionality of the “**shared genes**”, KEGG and GO analyses were performed. As the results show in figure 5a, the most significant GO biological process (BP) terms associated with the shared genes included cell adhesion, negative regulation of cell proliferation and nervous system development , while GO cellular component (CC) analysis revealed that the related proteins to these genes were mostly localized in plasma membrane and extracellular spaces and GO molecular function (MF) analysis showed that the related proteins functions to be calcium ion binding, extracellular matrix structural constituent, heparin, integrin and collagen binding. According to KEGG functional enrichment analysis, the shared genes participated in focal adhesion, calcium and cAMP signaling pathways (Figure 5b).

**Figure 5.**
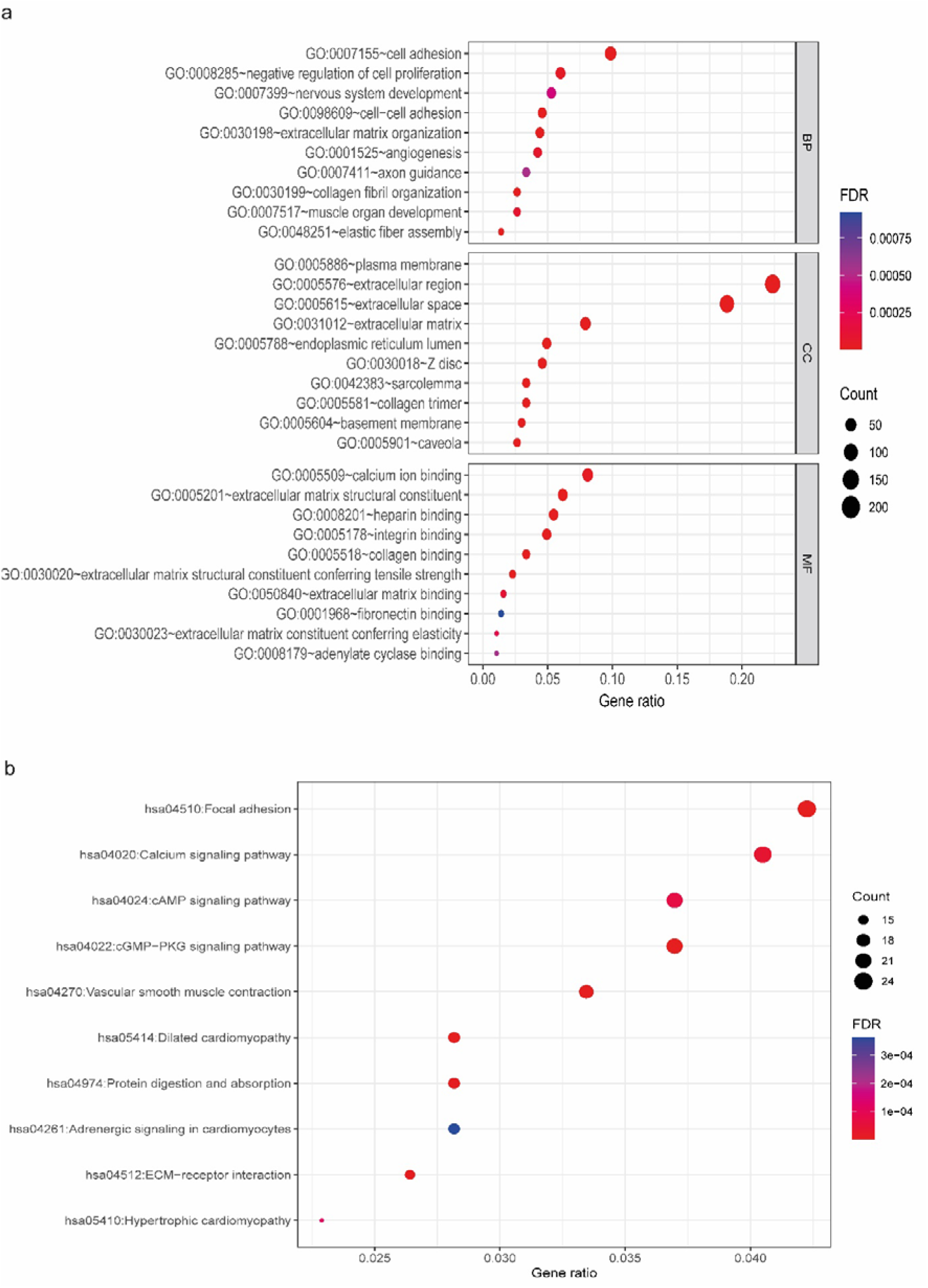
Dot plots of GO and KEGG functional enrichment analysis based on the shared genes. The horizontal and vertical axes represent gene ratio and terms, respectively. For each term, FDR<0.05 was considered significant. Top ten terms in each plot are presented. GO terms (a), KEGG terms (b). BP: biological process, CC: Cellular component, DEGs: differentially expressed genes, GO: Gene Ontology, FDR: false discovery rate, KEGG: Kyoto Encyclopedia of Genes and Genomes, MF: Molecular function, STAD: Stomach adenocarcinoma, TCGA: The Cancer Genome Atlas

Next, to investigate the interactions between the proteins associated with the “**shared genes**”, a PPI network was constructed which included 538 nodes and 587 edges (Figure 6a). The highest node degree centrality was attributed to FN1 (Fibronectin 1), followed by COL1A1 (Collagen Type I Alpha 1 Chain), COL3A1 (collagen type III alpha 1 chain), MMP2 (matrix metallopeptidase 2), COL1A2 (collagen type I alpha 2 chain), DCN (decorin), COL5A1 (collagen type V alpha 1 chain), THBS1 (thrombospondin 1), LUM (lumican), COL6A2 (collagen type VI alpha 2 chain) (Figure 6b).

**Figure 6.**
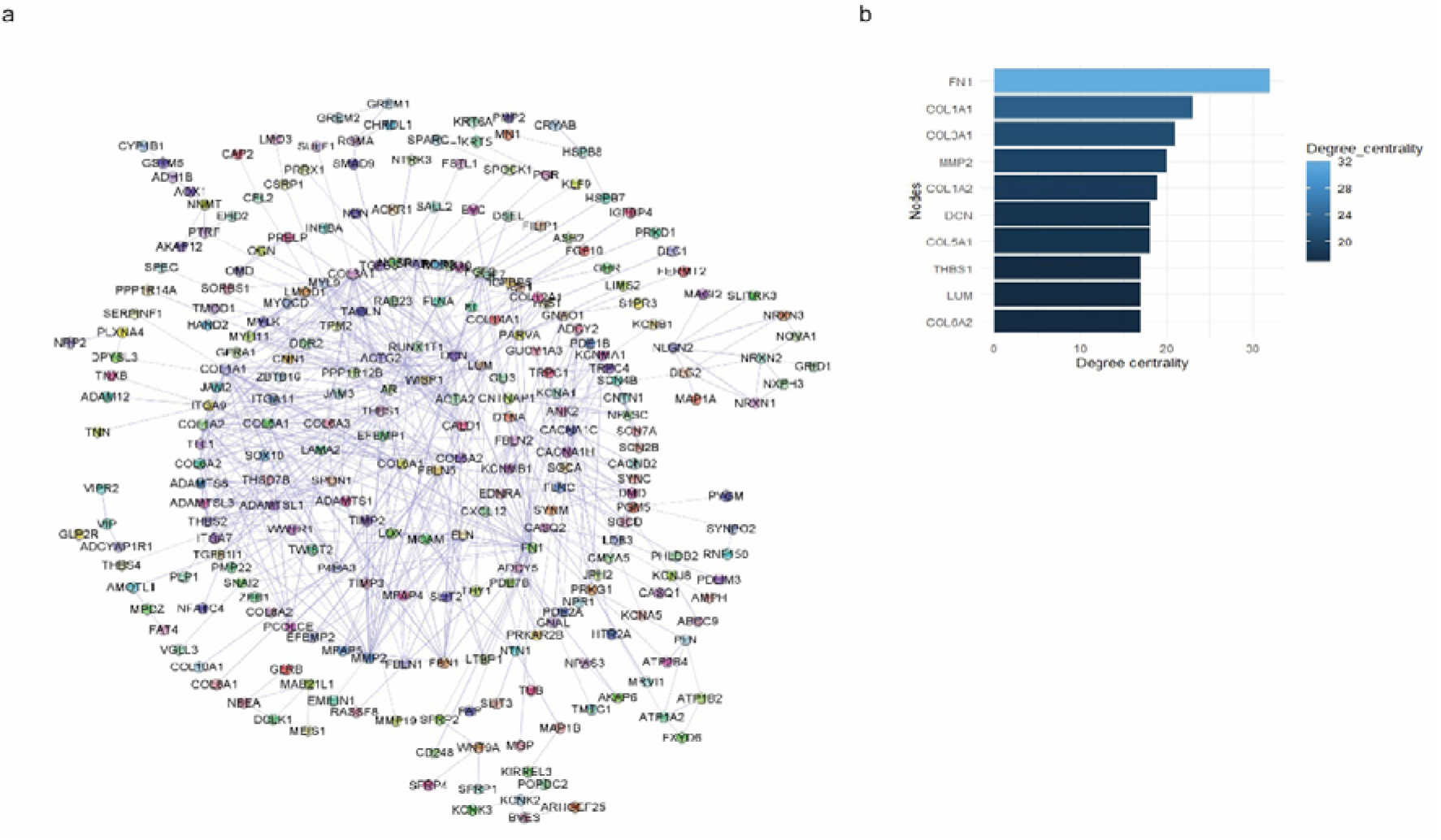
PPI analysis of the shared genes. The PPI network (a) and the bar plot of the top 10 nodes with the highest degree-centralities (b) are represented. PPI: Protein-protein interaction

### Survival analysis of the shared genes and construction of a decision tree

The association of the 569 “**shared genes**” with TCGA-STAD overall survival was investigated and 62 genes were found to be significantly (*P*. value < 0.05) associated with overall survival (Suppl table 1). Next, the survival-related genes were submitted for Recursive Partitioning to construct a prognostic decision tree model (Figure 7a). The 375 tumoral samples of the TCGA-STAD dataset were partitioned into training (0.75) and test (0.25) sets. The decision tree was constructed using eight genes, including *AKAP6*, *GLRB*, *LINC00578*, *LINC00968*, *MIR145*, *NBEA*, *NEGR1* and *RUNX1T1*. The efficiency of the model was evaluated using the ROC curves, according to which the AUC of the training and the test sets were 0.81 and 0.67, respectively (Figures 7b and c).

**Figure 7.**
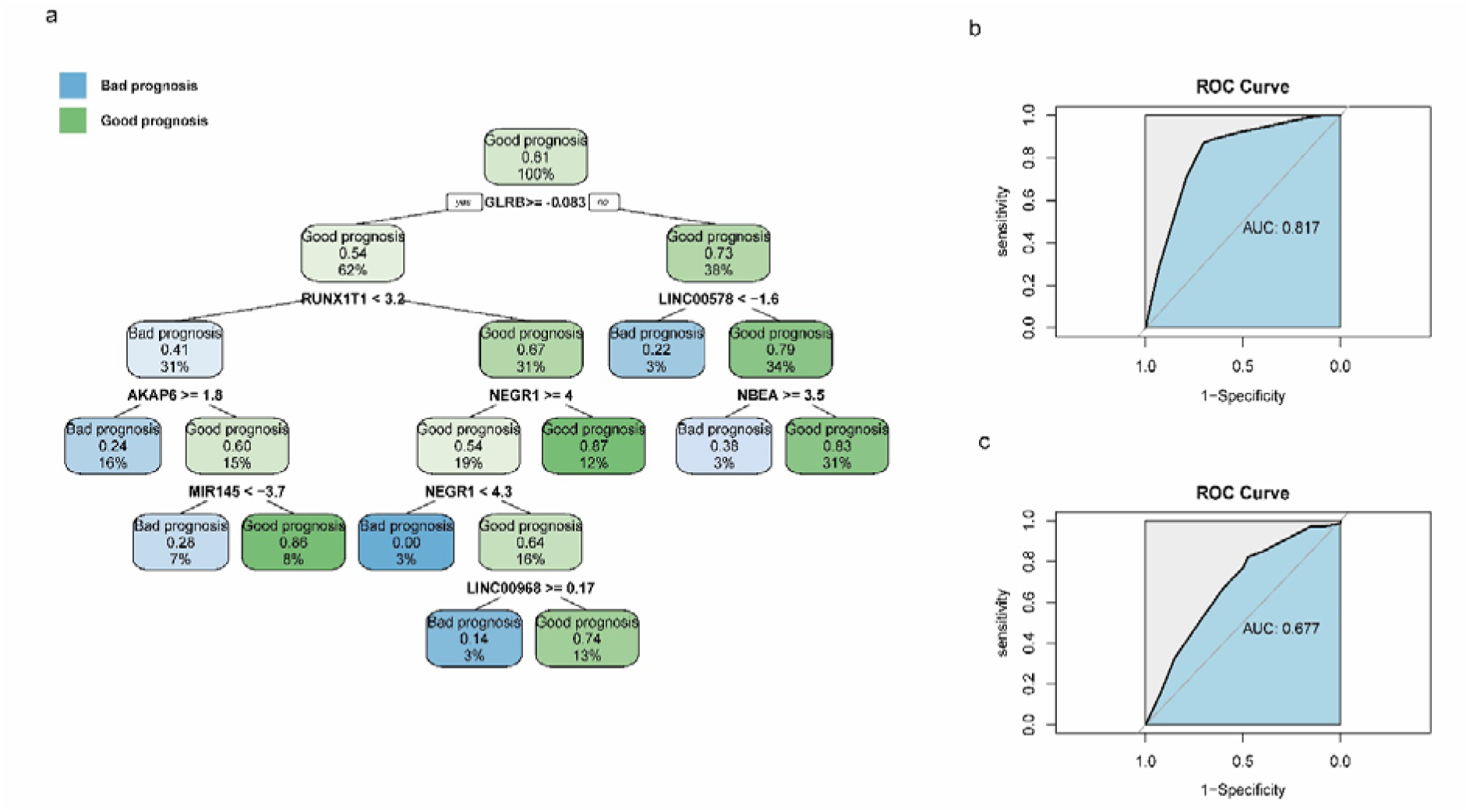
The prognostic decision tree model for GC constructed using the shared genes. The prognostic decision tree (a) and ROC curves of the training (b) and test (c) sets are represented. GC: Gastric cancer, ROC: Receiver operating characteristic

## Discussion

GC is one of the leading cancer-related mortalities worldwide. One of the characteristics of solid tumors is the hypoxic microenvironment, which by itself can lead to high stemness in the tumor cells (30). High stemness can result in de-differentiation of the tumor cells and increase the chances of invasion and metastasis (31). The resulting aggressive phenotype is associated with poor prognosis, and that’s the reason behind many studies’ interest in finding novel hypoxia- and stemness-related prognostic biomarkers (32–35). There are studies that have examined the combined effects of hypoxia and stemness on tumor progression (36,37) and in this regard, there is a lot to be discovered in the field of gastric cancer.

In the present study, the RNA-seq data of gastric cancer were retrieved from TCGA database and hypoxia and stemness scores were calculated. Hierarchical clustering and survival analysis were then carried out resulting in two clusters (one with low hypoxia and high stemness (cluster 1) and another one with both high hypoxia and high stemness (cluster 4)) showing the most difference in survival rates. Of note, performing the same pipeline on an external validation set (GSE62254) resulted in the same pattern.

Hypoxia within tumors has been linked to the decreased disease-free survival (DFS) outcomes across a variety of cancer types (38–42). In the same way, in the context of gastric cancer, several studies have reported the association between hypoxia and hypoxia-related gene expressions and poorer overall survival (43–45).

The hypoxic environment has the potential to alter the gene expression levels that regulate various metabolic and other physiological processes. Furthermore, the hypoxic signaling pathways interact with other cellular pathways to modify the malignant behavior of cancer cells including the proliferation, migration, invasion, and angiogenesis and significantly influences the treatment outcomes of cancer (46,47). The current study utilized TCGA data to evaluate hypoxia scores for gastric cancer and assess their correlation with survival rates. Our findings align with prior studies, demonstrating a direct correlation between elevated hypoxia levels and diminished survival rates.

The concept of “stemness” encompasses the coordinated molecular mechanisms that regulate and sustain the characteristics of stem cells (48). Since its introduction by Malta et al. (19), the mRNAsi score, derived from mRNA expression profiles, has become a widely utilized metric in numerous studies to evaluate the stemness properties of tumor specimens. Its application reveals divergent relationships with survival outcomes across distinct cancer types. While some malignancies like breast and hepatocellular carcinoma demonstrate an inverse correlation between mRNAsi levels and overall survival rates (49,50), indicating that higher mRNAsi scores are associated with poorer prognoses, others like lung adenocarcinoma, gastric and bladder cancers exhibit a positive correlation, suggesting that elevated mRNAsi scores correspond to improved overall survival rates (36,51,52). Moreover, in certain instances, no statistically significant associations between mRNAsi scores and survival outcomes are discerned (53,54). These findings suggest that the relationship between mRNAsi scores and survival outcomes may be tissue-dependent, varying significantly across different cancer types.

Of note, in the current study, we did not investigate the association between the mRNAsi index and survival outcomes in TCGA-STAD samples in isolation. Instead, we combined hypoxia and mRNAsi scores to assess their overall association with overall survival. Based on the findings of the present study, samples with high hypoxia and high stemness showed the worse survival rates compared to ones with low hypoxia and high stemness. Our analysis of the TCGA-STAD as well as an external GC dataset indicated that hypoxia is the most critical factor in determining the survival status of GC patients, whereas stemness appears to have a minimal impact on survival.

Differential expression analysis between clusters 1 (higher survival rates) and 4 (lower survival rates) of the TCGA-STAD samples resulted in 1446 genes. Performing WGCNA on the normalized RNA-seq data of TCGA-STAD resulted in 1885 hub genes showing associations with selected clinical traits. GO-BP analysis on the shared genes between these two gene sets showed an enrichment in cell adhesion and proliferation as well as nervous system development. Previous studies have shown that the nervous system plays a crucial role in promoting tumor metastasis by regulating metastatic cascades through the secretion of neural-related factors from nerve endings, such as neurotrophins, neurotransmitters, and neuropeptides(55). KEGG functional analysis also revealed that the shared genes were mostly involved in focal adhesion, calcium and cAMP signaling pathways, all of which are involved in cancer progression and invasion (56–58).

After performing survival analysis on the shared genes, a decision tree was constructed with eight genes including *AKAP6*, *GLRB*, *LINC00578*, *LINC00968*, *MIR145*, *NBEA*, *NEGR1* and *RUNX1T1*. AKAP6 is a member of the family of A-kinase anchoring proteins (AKAPs). AKAPs act as scaffolds that localize protein kinase A (PKA) to its target proteins, thereby modulating and enhancing the biological outcomes of cAMP signaling. cAMP–PKA signaling is a key pathway that influences cancer cell proliferation, motility, invasion and metabolism (58,59). Several studies have revealed the link between AKAP6 polymorphisms and cancer. For instance, it has been demonstrated that AKAP6 variants can influence the risk and outcome of glioma (60). Another study reported that AKAP6 mutations are increased in gastric cancer (61). Further research is needed to elucidate the mechanisms by which AKAP6 dysregulation impacts cancer progression. *GLRB* encodes the beta subunit of the glycine receptor chloride channel (GlyR β), which has a significant role in inhibitory neurotransmission in central nervous system and also modulates neurotransmitter release presynaptically (62). Studies have elucidated the importance of GLRB in cancers. For example, the widespread transcription of GlyR β observed in SCLC cell lines suggests its potential as a valuable marker for neoplastic cells (63). GLRB is also considered as a potential drug target for colorectal cancer (64). *LINC00578*, a recently identified long noncoding RNA (lncRNA), has been linked to prognosis in lung, breast, and pancreatic cancers (65–67). Furthermore, research suggests that *LINC00578* promotes the development and growth of pancreatic cancer by inhibiting ferroptosis, a type of cell death induced by iron-dependent oxidation (67). Long intergenic non-protein coding RNA 968 (*LINC00968*) may function as an oncogene in certain cancers, such as osteosarcoma (68), while acting as a tumor suppressor in others, such as lung adenocarcinoma (69). Further research is needed to elucidate the mechanisms by which *LINC00968* influences cancer progression. miR-145 plays a critical role in regulating gene expression and is notably downregulated in gastric cancer cells. Restoration of miR-145 activity has been linked to reduced cell proliferation and migration, alongside an increase in apoptosis in gastric cancer cells. These findings suggest that miR-145 holds significant potential as a therapeutic target for treating gastric cancer (70,71). Nbea, encoded by the *NBEA* gene, belongs to a large and diverse group of A-kinase anchor proteins that direct protein kinase A activity to specific subcellular locations by binding to its type II regulatory subunits. Nbea is involved in cargo transport and endomembrane compartmentalization, as well as in the formation of electrical and chemical synapses (72). Research has demonstrated an increase in NBEA mutations in gastric cancer (61), and its overexpression has been associated with a poorer prognosis in gastric cancer patients (73). Negr1 is a protein that belongs to the immunoglobulin LON family and functions as a GPI-anchored cell adhesion molecule. It plays a critical role in regulating synapse formation and neurite outgrowth (74). Analysis of gene expression profiles from tumor biopsies and normal tissues revealed that *NEGR1* is commonly downregulated in various human cancer tissues. Overexpression of *NEGR1* has been shown to suppress proliferation, anchorage-independent growth, and migration of human ovarian cancer cells (75). Conversely, another study identified Negr1 as a target of TGFβ, essential for maintaining the tumorigenic activity of metastatic breast cancer cells (76). Given the controversial results regarding the role of *NEGR1* in cancer, further research is necessary to elucidate its precise function and therapeutic potential in cancer progression. RUNX1T1 functions as a transcriptional repressor and is commonly regarded as a biomarker for primary pancreatic endocrine tumors, breast cancer, and colorectal cancer, as well as being a reliable predictor of patient prognosis (77). It is reported that *RUNX1T1* promoter is hypermethylated in GC, and its ectopic expression in GC inhibits cancer cells’ proliferation and overall is considered a tumor suppressor in GC (78).

This study highlights several promising areas for future research. Firstly, prospective studies could offer more control over sampling and data quality, addressing the limitations inherent in using retrospective data like TCGA. Additionally, given the heterogeneous nature of gastric cancer, future research should aim to minimize sampling bias by incorporating a diverse array of samples. Furthermore, the functionality of the proposed decision tree model warrants assessment on larger datasets, and further experimental analyses are necessary to evaluate the model’s efficiency.

## Conclusion

In conclusion, this study integrates bioinformatics approaches to elucidate the interplay between hypoxia and stemness in gastric cancer and its implications for patient prognosis. By analyzing TCGA data and an external validation dataset, the study shows that combining hypoxia and stemness scores can categorize GC patients into different prognostic groups. Specifically, clusters characterized by high hypoxia and stemness exhibit poorer survival outcomes compared to those with low hypoxia and high stemness. Differentially expressed genes identified between these clusters, along with hub genes derived from WGCNA, highlight pathways related to cell adhesion, proliferation, and nervous system development, emphasizing their role in GC progression. Furthermore, the construction of a prognostic decision tree model based on survival-associated genes provides a framework for predicting patient outcomes. This approach highlights the potential of hypoxia and stemness as biomarkers for GC prognosis and suggests further validation and therapeutic strategies for managing this cancer.

## Statements & Declarations

### Author Contributions

Sharareh Mahmoudian-Hamedani: Data Curation (lead); Formal Analysis (equal); Investigation (equal); Methodology (equal); Software (lead); Visualization (lead); Writing – Original Draft Preparation (equal); Writing – Review & Editing (equal)

Maryam Lotfi-Shahreza: Formal analysis (equal); methodology (equal); validation (equal); writing – review and editing (equal)

Parvaneh Nikpour: Conceptualization (lead); Funding Acquisition (lead); Investigation (equal); Project Administration (lead); Resources (lead); Supervision (lead); Validation (lead); Writing – Original Draft Preparation (equal); Writing – Review & Editing (equal)

### Data Availability

The results published or shown here are in whole or part based upon data generated by the TCGA Research Network (https://portal.gdc.cancer.gov/legacy-archive/search/f) and GEO database (https://www.ncbi.nlm.nih.gov/geo/). Data are available upon request to the corresponding author.

